# Parasite genotype is a major predictor of mortality from visceral leishmaniasis

**DOI:** 10.1101/2022.07.28.501951

**Authors:** Cooper Alastair Grace, Kátia Silene Sousa Carvalho, Mayara Ingrid Sousa Lima, Vladimir Costa Silva, João Luís Reis-Cunha, Matthew J. Brune, Sarah Forrester, Conceição de Maria Pedrozo e Silva de Azevedo, Dorcas Lamounier Costa, Doug Speed, Jeremy C. Mottram, Daniel C. Jeffares, Carlos H.N. Costa

**Affiliations:** York Biomedical Research Institute, Department of Biology, University of York, York YO10 5DD, United Kingdom; Department of Community Medicine and Institute of Tropical Diseases Natan Portela, Federal University of Piauí, Teresina, Brazil; Department of Biology, Postgraduate Programs in Health Sciences and Postgraduate Program in Health and Environment, Federal University of Maranhão, São Luís, Maranhão, Brazil; Department of Medicine and Postgraduate Program in Health Sciences, Federal University of Maranhão, São Luís, Maranhão, Brazil; Intelligence Center for Emerging and Neglected Diseases (CIATEN), Teresina, PI, Brazil; Mother Child Department, Federal University of Piauí, Teresina, PI, Brazil; Centre for Quantitative Genetics and Genomics, Aarhus University, Denmark

**Keywords:** Visceral leishmaniasis, Brazil, *Leishmania infantum*, virulence factors, genetic diversity, quantitative genetics, mortality

## Abstract

**Background:** Visceral leishmaniasis (VL) is a potentially fatal disease mainly caused by *Leishmania infantum* in South America and *L. donovani* in Asia and Africa. Disease outcomes have been associated with patient genotype, nutrition, age, sex, comorbidities, and co-infections. In this study, we examine the effects of parasite genetic variation on VL disease severity in Brazil.

**Methods:** We collected and sequenced the genomes of 109 *L. infantum* isolates from patients in northeast Brazil and retrieved matching patient clinical data from medical records, including mortality, sex, HIV co-infection and laboratory data (creatinine, haemoglobin, leukocyte and platelet counts). We identified genetic differences between parasite isolates, including single nucleotide polymorphisms (SNPs), small insertions/deletions (indels), and variations in genic, intergenic, and chromosome copy numbers (copy number variants, CNVs). To describe associations between the parasite genotypes and clinical outcomes, we applied quantitative genetics methods of heritability and genome-wide association studies (GWAS), treating clinical outcomes as traits that may be influenced by parasite genotype.

**Findings:** Multiple aspects of the genetic analysis indicate that parasite genotype affects clinical outcomes. We estimate that parasite genotype explains 83% chance of mortality (narrow sense heritability, *h*^2^ = 0·83±0·17), and has a significant relationship with patient sex (*h*^2^ = 0·60±0·27). Impacts of parasite genotype on other clinical traits are lower (*h*^2^ ≤0·34). GWAS analysis identified multiple parasite genetic loci that were significantly associated with clinical outcomes; 17 CNVs that were significantly associated with mortality, two with creatinine and one with bacterial co-infection, jaundice and HIV co-infection; and two SNPs/indels and six CNVs that associate with age, jaundice, HIV and bacterial co-infections, creatinine, and/or bleeding sites.

**Interpretation:** Parasite genotype is an important factor in VL disease severity in Brazil. Our analysis indicates that specific genetic differences between parasites act as virulence factors, enhancing risks of severe disease and mortality. More detailed understanding of these virulence factors could be exploited for novel therapies.

**Author Summary:** Multiple factors contribute to the risk of mortality from visceral leishmaniasis (VL), including, patient genotype, comorbidities, and nutrition. Many of these factors will be influenced by socio-economic biases ^1^. Our work suggests that the virulence of the infecting parasite is an important risk factor for mortality. We pinpoint some specific genomic markers that are associated with mortality, which can lead to a greater understanding of the molecular mechanisms that cause severe VL disease, to genetic markers for virulent parasites and to the development of drug and vaccine therapies.

## Introduction

Leishmaniasis is a neglected tropical disease caused by protozoan parasites of the *Leishmania* genus. Visceral leishmaniasis (VL), the most severe form, is caused by *Leishmania infantum* in South America, Mediterranean Basin, Middle-East and Central Asia and *L. donovani* in East Africa and the Indian Subcontinent. VL is characterised by fever, weight loss, hepatosplenomegaly, anaemia, pancytopenia, hypoalbuminemia, and hypergammaglobulinemia, and is frequently fatal if untreated.^2^ Severe disease presents as hepatic dysfunction with jaundice, oedema, and dyspnoea.^3^ Death is usually associated with hemorrhagic phenomena, which could be caused by bacterial co-infection^4,5^ as a consequence of cytokine driven systemic inflammation.^6^ *Leishmania*/HIV results in greater risks of relapse and fatality.^7,8^ Despite the range of clinical presentations, the parasite factors that determine differential virulence in humans are not well understood.

There are approximately 5000 VL cases/year in Brazil, 90% of the cases registered in the Americas.^9–12^ There are large numbers of asymptomatic *L. infantum* infections, with various studies identifying 1-6% of the general population in endemic regions.^7,13,14^ VL case fatality rates in Brazil average 7%, the highest in the world, and show an increasing trend.^12^ Hence, understanding the factors that lead to severe disease and mortality is a priority for leishmaniasis research. Disease outcomes are known to be associated with patient age, sex, comorbidities,^4^ adverse effects of treatment, and co-infections such as HIV.^15^ Genome-scale association studies (GWAS) have shown to human genotypes influence VL disease severity in East Africa, Brazil and the Indian subcontinent,^16^ including a strong influence from the HLA-DR-DQ region within the major histocompatibility complex.^17^ However, apart from studies of drug-resistance,^10,18^ the influence of natural parasite genetic variation on disease severity has not been investigated using genome-scale methods.

To investigate this, we collected 109 *L. infantum* isolates from Piauí and Maranhão states in Brazil that have large VL burdens.^19,20^ We use genome-scale quantitative genetic analyses to investigate the effects of parasite genotype on multiple clinical indicators of disease severity and mortality. Our analysis showed that parasite genotype influences multiple aspects of disease severity and a strong relationship with patient mortality. We identify multiple parasite genetic loci that affect mortality. These virulence factors are key points for vaccine and drug discovery,^21^ that will be complementary to reverse vaccinology methods.^22^

## Methods

### Study design and sample collection

The objective of this study was to search for relationships between parasite genotypes and patient clinical factors and outcomes, including mortality. To gather parasite genotypes, we collected *L. infantum* parasite samples from 109 diagnostic aspirates of 109 patients in Piauí and Maranhão states (**Figure 1**), cultured them and sequenced their genomes. DNA extraction from promastigote parasites was performed after two culture passages, to minimise the impact of culture adaptation. Each parasite genome is matched to clinical patient data. The corresponding patient and clinical data was extracted from medical records and routine patient follow-up. This included patient factors (age, sex), clinical characteristics (mortality, bleeding, oedema, HIV coinfection, jaundice, vomiting, dyspnoea, bacterial co-infection), and laboratory clinical metrics (leukocyte and platelet counts; creatinine and haemoglobin levels). We also estimated probabilities of mortality from clinical models (Kala-Cal*, KalaCal**) (**Supplementary Table S1; S2**)^5^. Our data analysis employed quantitative genetics methods of narrow sense (additive) heritability and genome-wide association studies (GWAS) to associate specific parasite genetic variants (SNPs, indels, gene and chromosomal CNVs) with patient factors and clinical characteristics.

**Figure 1.**
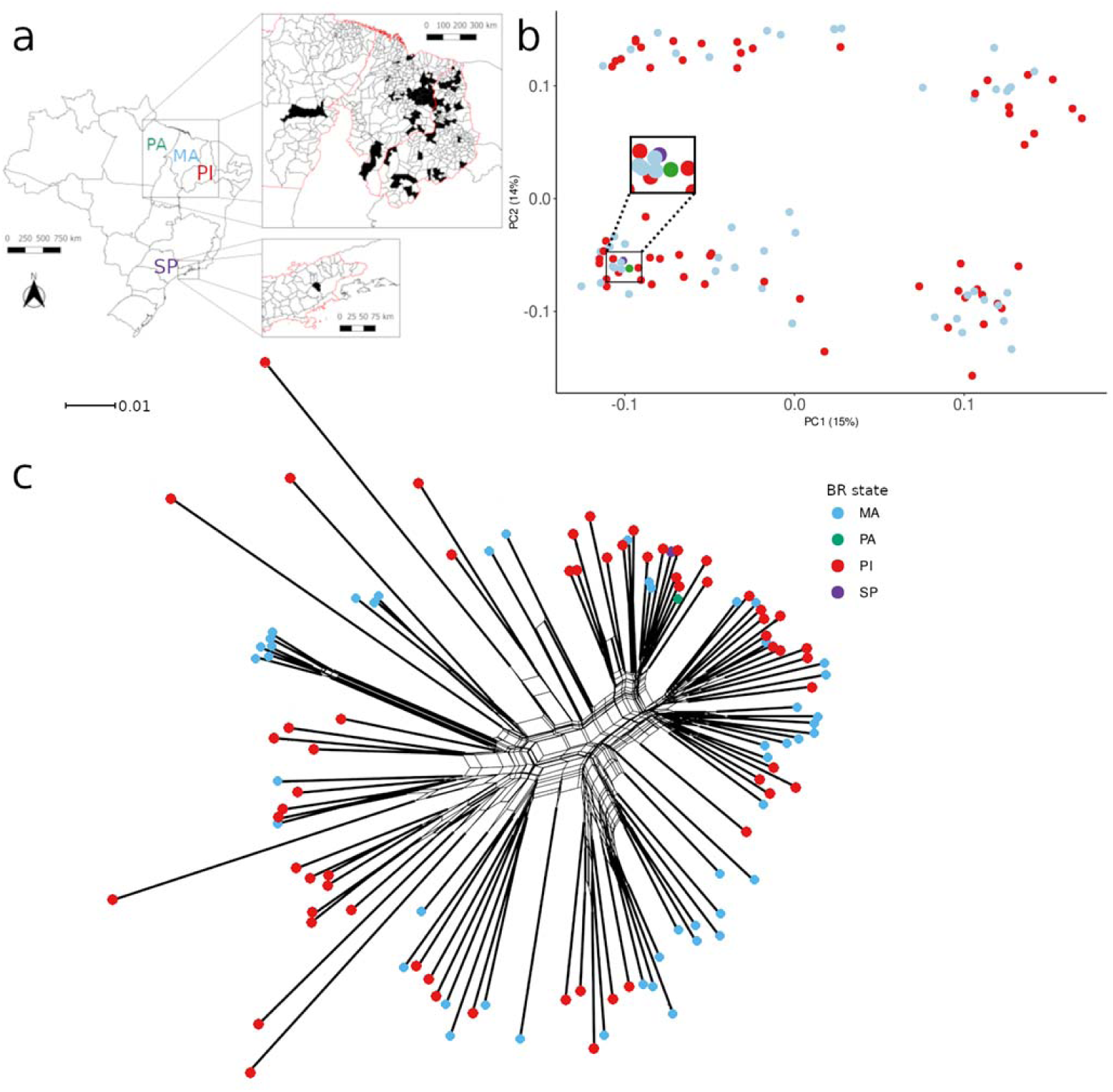
Sample origins and population structure. **Panel A**: Map showing residential locations of VL patients that provided the *L. infantum* strains. States in Brazil are: MA - Maranhão; PI - Piauí; PA - Pará; SP - São Paulo. States Pará and São Paulo possess a single isolate each. **Panel B**: principal component coordinates, derived from SNP variation, shows that genetic relatedness is not clustered by state. Each filled circle indicates the principal-component derived relatedness between isolates. Circles are coloured by their state of origin. **Panel C**: Phylogenetic network analysis of all 109 strains, with coloured circles indicating state of origin using the same colour scheme as **Panel B**. Branches with thick lines have placement with 100% bootstrap support. Scale bar indicates the number of substitutions per site (network file and version with isolate names are available in the FigShare depository).

### Ethics

Piauí samples and patient data were obtained as a part of a broad study from the Federal University of Piauí and Federal University of Maranhão, respectively. Patient recruitment was performed at the Institute of Tropical Medicine Natan Portella, approved by the Research Ethics Committee of the Federal University of Piauí (approval ID number 0116/2005); and in the Reference Hospital for Infectious Diseases of the Maranhão, approved by the Research Ethics Committee of the University Hospital of the Federal University of Maranhao (approval ID 2.793.599) and the Research Ethics Committee of the Federal University of Maranhao (approval ID 3.921.086). Both projects were approved by the Department of Biology Ethics Committee, at the University of York (approval IDs DJ201803, DJ202009). All methods were performed according to the approved guidelines and regulations. All participants or their legal guardians signed a written informed consent form. Clinical, epidemiological and laboratory data from patients were obtained during hospital admission, medical follow-up or medical records. All patients were treated with one of the following protocols, based on disease severity: 1. Pentavalent antimony (20mg/kg/day, maximum 800mg/day) for 21 days; 2. Amphotericin B deoxycholate 1mg/kg/day for 14-20 days); or 3. Ambisome (Liposomal Amphotericin B, 3mg/kg/day for 7 days for immunocompetent and 14 days for immunocompromised patients). Parasites were isolated from bone marrow aspirates before starting patient treatment, and culturing was performed at the Laboratory of Leishmaniasis at the Universidade Federal do Piauí, or at the Laboratory of Genetics and Molecular Biology at the Universidade Federal do Maranhão.

### Sequencing and bioinformatic analyses

Genome sequencing was performed on Illumina HiSeq 2500, generating paired-end 150nt reads. All but eight of the libraries had genome coverage ≥30x (median 49x). (**Supplementary Table S1**). Sequencing of 73 isolates has previously been described.^23^ Sequencing reads were mapped onto *L. infantum* JPCM5 v.46 (https://tritrypdb.org) reference genome using bwa v.0.7.17.^24^ Read duplicates were removed with SAMtools v.1.9.^25^ For each strain, SNPs and indels were called using Genome Analysis Toolkit (GATK) *HaplotypeCaller* v.4.1.0.0^26^ and the ‘discovery’ genotyping mode of Freebayes v.1.3.2 (https://github.com/ekg/freebayes), only accepting calls discovered by both methods. We retained only biallelic variants, with read depth within 0·3-1·7x the chromosome coverage, excluding any variants called on repetitive regions (see **Supplementary Methods** available on FigShare). All genomes were initially compared using phylogenetic approaches and Principal Component Analysis (PCA; see **Supplementary Methods**). Alignments and resulting phylogenetic tree files are available on FigShare. The gene, intergenic, and chromosome copy number variation of the 109 isolates was also assessed. This was performed by comparing variations in read depth coverage (RDC) between isolates (see **Supplementary Methods**).

### Genetic analysis; estimating heritability and genome-wide association studies (GWAS)

SNPs/indels, genes, intergenic region, and chromosome CNVs were used as genotypes to estimate the total effect of parasite genetic variation on the patient clinical traits (narrow sense heritability), using LDAK v.5.1.^27^ In this analysis, clinical data were coded either as binary or quantitative traits (**Supplementary Table S2**). PLINK v.1.9^28^ was used to create the binary files. Gene, intergenic, and chromosome CNVs were generated from RDC values normalised by genome coverage for each gene, intergenic region, or chromosome and coded as quantitative genotypes. Gene CNVs varied from 0 to 3·8 copies per haploid genome copy (mean 1·017). The contribution of genotype class to each clinical trait was estimated using LDAK restricted maximum likelihood (REML; see **Supplementary methods**).

We used GWAS to identify specific genomic variants that were statistically associated with each clinical trait. LDAK --linear was used to conduct mixed model GWAS, using the kinship matrix derived from SNP/indel variants to control for unequal strain relatedness. Significance thresholds were determined via trait permutation. For each trait, values were permuted 1000 times, and parasite genotype-human trait associations estimated. The lowest *P*-value was determined from each replicate permutation analysis, and the 5^th^ percentile used as the significance threshold for associations (i.e. 5% family-wise false discovery rate) for each trait. *P*- value thresholds varied from 5·2 to 7·3 (-log10). GWAS for SNPs/indels used only biallelic sites with call quality ≥2000 and population-wide MAF >5% across all 109 isolates (ie: an allele must be present in at least 11 of the 218 haploid chromosomes to be used; *n*=3,526). GWAS for CNVs used only genes with the mean coverage >20% of genome coverage and coverage coefficient of variation (CV) (SD/mean) >1% (*n*=8,219 genes; see **Supplementary methods**). Scripts used to run associations and permutations as well as the script to generate the Manhattan, Q-Q, correlation, and boxplots are available on Figshare (URL to be provided upon manuscript publication).

### Role of the funding source

The funders had no role in study design, patient recruitment, collection, analysis, or interpretation of data, in the writing of the manuscript or in the decision to submit for publication.

## Results

### Sampling and clinical traits

We collected *L. infantum* isolates for genomic sequencing and clinical data from 109 patients with VL in Brazil. The isolates were collected from patients in the highly endemic Piauí and Maranhão states (**Figure 1, Panel A**). Each parasite isolate was matched to the source patient with information on age, sex, binary clinical factors mortality (recovered/died), bleeding, presence of oedema, HIV status, jaundice, vomiting, dyspnoea, bacterial co-infection; laboratory clinical metrics (leukocyte and platelet counts; haemoglobin and creatinine levels); and estimated probabilities of mortality from clinical models (Kala-Cal*, Kala-Cal**).^5^ All these factors varied among patients (**Supplementary Figure 13, Supplementary Tables S2**; **S3** available on FigShare), and mortality was well-predicted by the Kala-Cal* model (**Supplementary Figure 14**). In this cohort, no trait, except age, was associated with HIV status (**Supplementary Figure 15**). The only other between-trait correlations were that leukocyte and creatinine counts declined with patient age (**Supplementary Figure 16**).

### Genomic variation

To characterise genomic variation in these 109 *L. infantum* isolates, we identified SNP/indel variants, gene (GCNVs), intergenic (IGCNVs), and chromosome copy variations (CCNVs) (**Table 1**). In accordance with other analyses,^19,23,29^ *L. infantum* populations contain tenfold less genetic diversity than *L. donovani* populations in East Africa (nucleotide diversity in Brazil π=4·3×10^−5^, Africa π=42×10^−5^), and half as much diversity as the Indian subcontinent (India π=8×10^−5^). Analysis of between-strain relatedness with principal components analysis (PCA) and phylogenetics showed very little population structuring, with no clear distinction between isolates based on state of origin (**Figure 1, Panels B-C**). The majority of chromosomes were disomic (**Supplementary Figure 2**), whereas a high number of chromosomal duplications and gene CNVs were observed in several isolates, as expected for *Leishmania* and other eukaryotic genomes.^10,30^

**Table 1.** *L. infantum* genomic variability of 109 isolates.

| Genomic variation | Count |
| --- | --- |
| SNVs | 14,203 |
| SNPs | 4,468 |
| indels | 9,735 |
| variant sites MAF >5% | 3,526 |
| biallelic variants | 3,004 |
| genes | 8,748 |
| genes with CoV >1% and coverage >20% of genome coverage | 8,218 |
| intergenic regions | 8777 |
| intergenic regions with CoV >1% and coverage >20% of genome coverage | 8,219 |

Notes: SNVs - small nucleotide variants; MAF - minor allele frequency; CoV - coefficient of variation

### Effects of genetic variation in *Leishmania infantum* on clinical factors

To quantify the effect of parasite genetic variation, we estimated the narrow sense (additive) heritability, using clinical factors as traits (**Supplementary Table S2**) and parasite genetic variants as genotypes (see Methods). This analysis estimates the proportion of variation in patient clinical factors that is attributed to parasite genetic variation. We utilised four categories of genetic variants in this analysis; 1) SNP/indel variants, 2) chromosome copy variation (CCNV), 3) gene copy number variation (GCNV) and 4) intergenic region copy variation (IGCNV). Heritability estimation was performed using each genotype category independently, followed by a composite model that estimates the combined contribution of all variants. As we sought to examine the effects of paraste variation, host genotypes were not considered in this analysis.

Our results show that parasite genotype has significant effects on multiple clinical factors (**Figure 2**). Due to the small sample size (*n*=109), all estimates have relatively large standard deviations (**Supplementary Table S4**). Mortality was strongly affected by parasite genetic variation, with total heritability 0·83 (± standard deviation 0·17, 95% confidence interval 0·66-1·00), impacted by SNPs/indels and GCNVs. Sex was also strongly associated with SNP/indel variation (0·60±0·27, 95% confidence interval 0·33-0·87), indicating that parasite genotype influences the susceptibility of males and females to VL differentially. Strain-specific differences that affect male and female disease severity may be the cause of the higher occurrence of infections in males.^15^ The majority of the other traits were affected primarily by SNP/indel variations, rather than CNVs. Approximately a third of clinical traits (*n*=5) are affected by independent heritability components, while the majority of traits (*n*=11) are complex, with multiple types of genotypic variation contributing to host phenotype. This analysis shows that even within Brazil where *L. infantum* genetic diversity is relatively low, differences between parasites influence disease progression.

**Figure 2.**
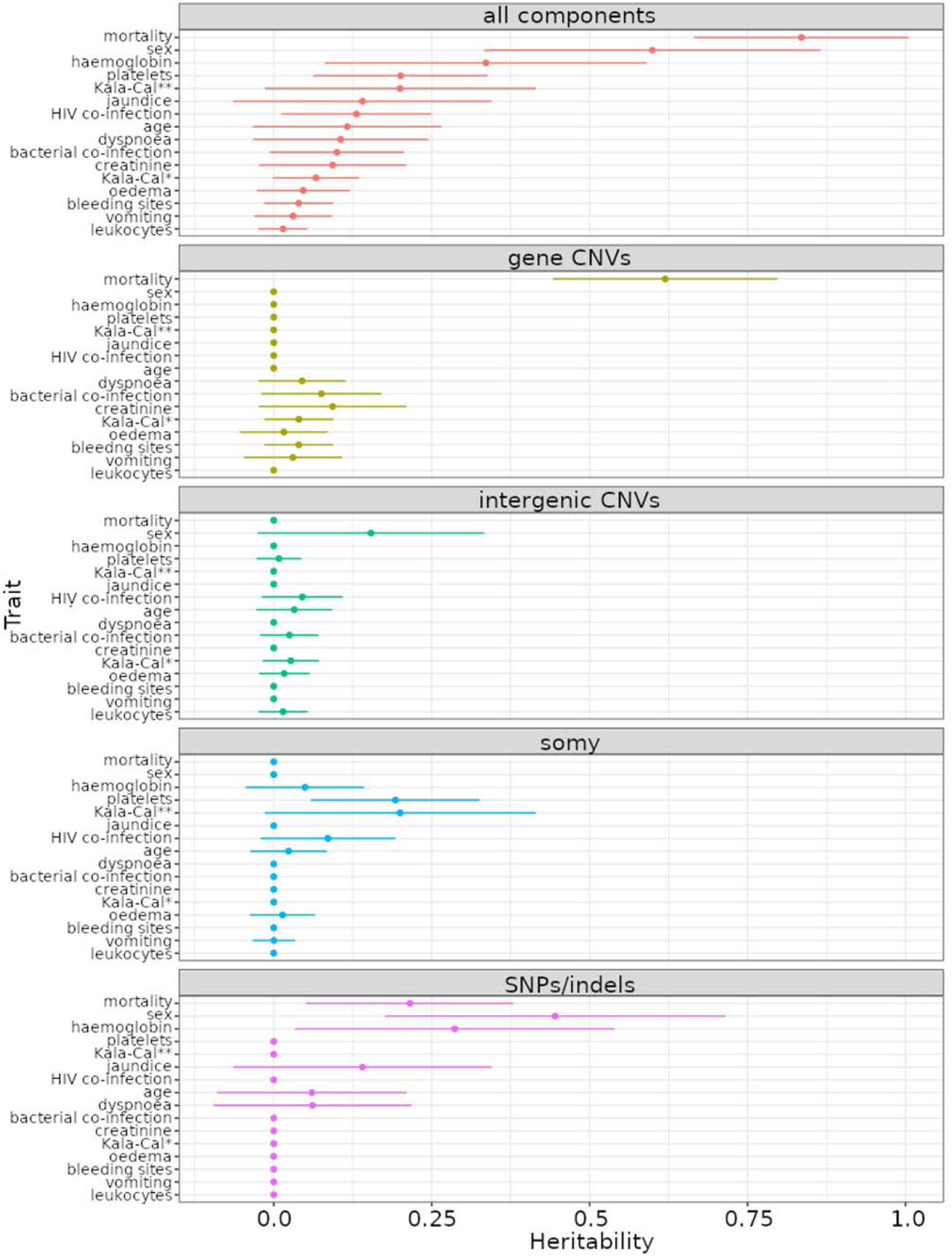
Heritability estimates from the composite model. Heritability estimates used normalised trait values. Horizontal lines represent standard deviations for each heritability estimate. Here most analyses used *n*=109 samples. The topmost panel shows the estimate of total heritability from all types of genetic variants combined. Successive panels below this show the heritability contributed by the various types of genetic variants. Due to linkage between different categories of variants (SNPs, gene copy variants etc.), the total heritability explained is not expected to be the sum of all variant types.

The effect of parasite genotype on mortality could be due to one lineage of particularly virulent parasites. To assess this possibility, we visualised this clinical outcome on a phylogenetic tree of isolates (**Supplementary Figure 4**). Isolates from patients who died are dispersed throughout the phylogeny, indicating that no specific lineage is responsible for mortality. This is expected, if multiple loci affecting mortality were segregating within the population and these loci are unlinked by recombination between strains. In this scenario, no single phylogeny will cluster these unlinked variants.

To identify the genetic loci that affect clinical outcomes, we conducted GWAS, using all four categories of genetic variants as predictors of clinical traits. In all analyses apart from chromosome copies (CCNVs), association *P*-values were inflated above expected values, indicating that multiple genetic loci affect clinical outcomes (**QQ plots, Supplementary Figures 5-12**). We observed no association between CCNV and any of the evaluated traits, which is in accordance with somy variation being only associated with short-term environmental adaptation^31^ that may vary after culturing.^32^ Given the small sample size available (*n*=109), we expect to discover only cases where genotypes have strong effects on clinical outcomes. Nevertheless, we find multiple parasite genetic variants that were associated with clinical outcomes above our permutation-derived thresholds (**Table 2**; **Supplementary Tables S5**-**S6**; **S8**).

**Table 2.** Genomic variations associated with clinical traits.

| **GWAS component** | **No. of significant associations** | **Trait(s)** |
| --- | --- | --- |
| SNPs/indels | 2 |  |
| GCNVs | 11 |  |
| IGCNVs | 12 |  |
| CCNV | 0 | N/A |

Two small variants were significantly associated with clinical factors. An insertion/deletion (indel) variant was significantly associated with the presence of jaundice, and a SNP variant was associated with patient age (*P*=5·2×10^−7^ and *P*=2·6×10^−6^, **Supplementary Table S5; Supplementary Figure 5**). Both of these variants were located within intergenic regions (**Supplementary Text B**). Gene copy variants were significantly associated with three traits: mortality (**Figure 3**), creatinine levels, and jaundice (**Supplementary Figure 7**). Variations in intergenic DNA copies were associated with four clinical factors: bacterial co-infection, creatinine, mortality and HIV co-infection (**Supplementary Figures 8-9** and **Supplementary Figures 21-23** in **Supplementary Text C** available on FigShare).

**Figure 3.**
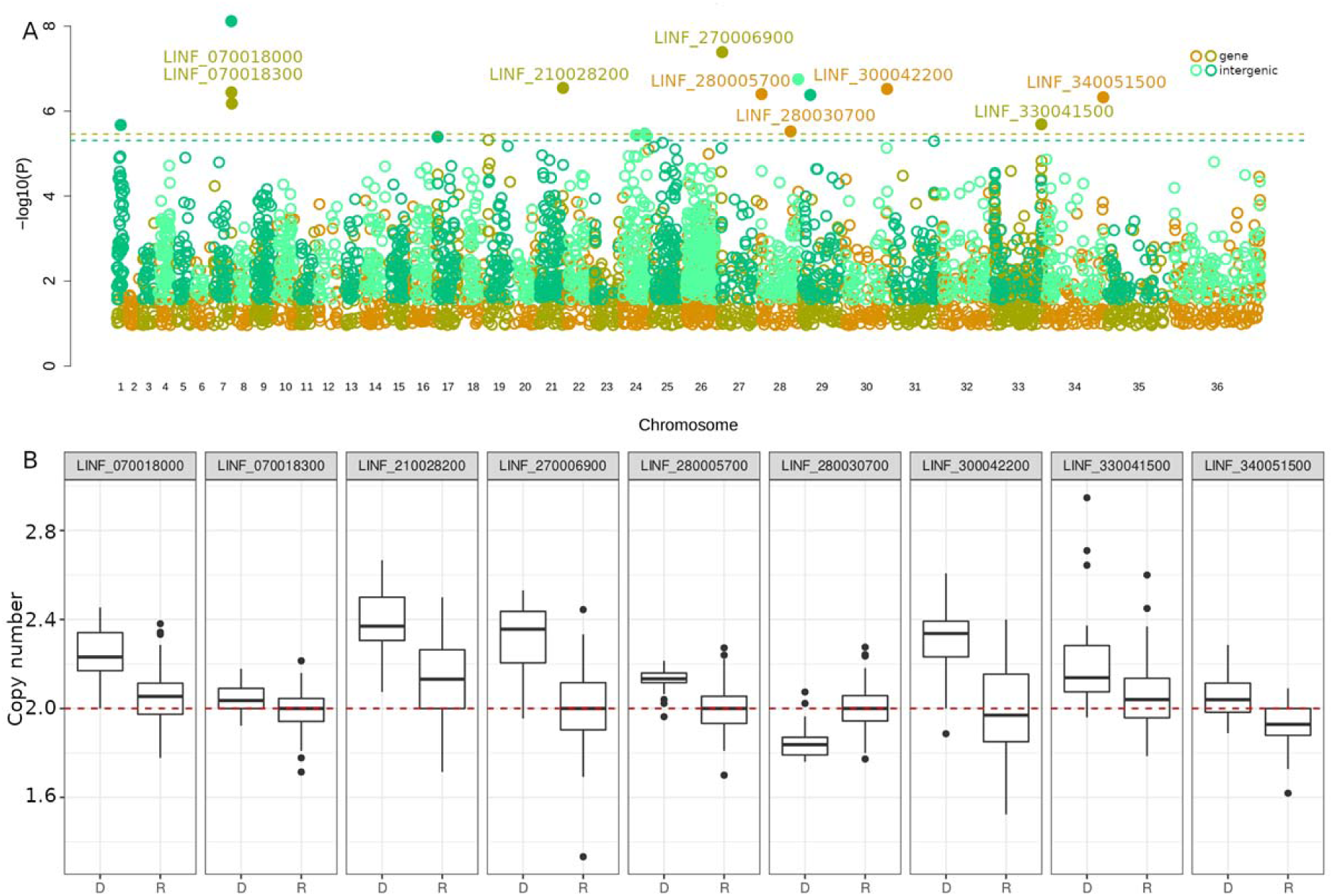
Association between genetic variation and mortality. **Panel A**: Manhattan plot displaying significant associations between G+CNVs (yellow/orange), and IGCNVs (dark/light green), and the mortality phenotype. Horizontal dashed lines indicate significance thresholds, calculated with 1000 permutations (see **Methods**). Significant variants are highlighted by filled circles. Significant GCNVs are also named. -log10 *P*-values were filtered to remove the 50% least significant values to aid figure clarity. **Panel B**: The copy numbers of significant GCNVs are displayed as boxplots, indicating copies from isolates where the patient either died (D) or recovered (R). The horizontal red dashed line indicates expected disomy. Mann-Whitney tests confirm that copy numbers for these genes are significantly different in parasites causing patient mortality: LINF_070018000, *P*=1·26×10^−7^; LINF_070018300, *P*=2·99×10^−3^; LINF_210028200, *P*=1·60×10^−8^; LINF_270006900, *P*=1·17×10^−8^; LINF_280005700, *P*=1·89×10^−8^; LINF_280030700, *P*=5·76×10^−8^ ; LINF_300042200, *P*=4·59×10^−8^; LINF_330041500, *P*=1·45×10^−3^; LINF_340051500, *P*=1·58×10^−7^.

The association scores, chromosomal position, and annotation of the 11 significant genes and 12 intergenic regions are displayed respectively in **Supplementary Tables S6** and **S8**. As expected, given the evidence for recombination in these isolates, we find that some strains contain multiple virulence-associated genotypes, and others few (**Supplementary Table S7**), which may result in a range of virulence levels.

The progression of disease is likely to be influenced by interactions between patient factors, comorbidities and parasite genotype. For example, it is possible that particular parasite genotypes are more likely to infect HIV positive patients that are immune-suppressed. To examine this possibility, we conducted analysis of gene and intergenic copy variants that affect mortality, using all other clinical traits as covariates. This analysis frequently produced lower (more significant) association *P*- values (**Supplementary Tables S9** and **S10**). The most significant association using covariates was between the gene LINF_340051500 (phosphatidylinositol 3-kinase *tor2*) copy variant and mortality (*P*=6·24×10^−9^), using Kala-Cal** risk prediction as a covariate. Since the Kala-Cal** mortality prediction uses many clinical characteristics, it likely captures many factors of the relationship between clinical factors, parasite and mortality. The gene LINF_270006900 (hypothetical protein) was still significantly associated with mortality when each other trait was individually used as a covariate and when all the other traits excluding Kala-Cal* and Kala-Cal** were simultaneously used as covariates (**Supplementary Table S9**). While these results do not define the precise relationships between parasite, host and factors, they show that genetic association studies have considerable promise.

## Discussion

We used genome data from 109 *L. infantum* isolates from northeast Brazil, coupled with detailed patient clinical data to study the effects of parasite variation on VL disease. We found that the chance of mortality and patient sex were strongly associated with parasite genotype (heritability 0·83±0·17 and 0·60±0·27, respectively). Although our sample size is very small, we find multiple associations that are supported by permutation-derived thresholds, for several clinical traits that are not correlated with one another. Both these significant GWAS associations (**Figure 3**) and the inflation of *P*-values in the SNP ‘QQ plots’ indicate that multiple genetic factors contribute to different aspects of clinical outcomes. While this sample size is very small, similarly small sample sizes have identified significant association with microbial GWAS.^33,34^

We expect that disease outcomes and clinical manifestations are a combination of the interaction between the parasite, patient, environmental factors, and treatment. While no study has investigated the effects of human genetic variation on disease severity or mortality, several studies have evaluated the effects of human genetic variation to both visceral and cutaneous leishmaniasis *susceptibility* using GWAS.^17^

The VL analysis identified a single locus with high significance; the Human Leukocyte Antigen (HLA) haplotypes in the HLA-DR-DQ region (*P* = 2.76 × 10^−17^) were the only genetic determinant of susceptibility to VL,^17^ that was consistent between populations in Brazil and India. Much weaker association hits (top hits at *P*<5×10^−5^), were observed for susceptibility to cutaneous leishmaniasis (CL). Our analysis of parasite genetic variants is entirely consistent with these results, if we consider that host-parasite interactions determine clinical severity. The most striking result of our analysis is the magnitude of the contribution from the parasite genotype that we detect, without considering the human genotype. We consider this result reasonable from two perspectives. Firstly, the effects of human genotypes of disease severity have not been examined directly (only disease susceptibility), and secondly, it is the *parasite* that causes disease, while host genotypes merely react to, and modify, disease severity.

Parasite-derived drug-resistance and drug pharmacokinetics are also important factors.^35^ Genome-scale approaches have identified loci in *L. donovani* that are associated with antimonial resistance, which affect clinical outcomes.^36,37^ Similarly, the low efficacy of miltefosine in Brazil compared to initial efficacies in the Indian subcontinent, has been associated with the deletion of the Miltefosine Sensitivity Locus (MSL) that has only been detected in *L. infantum* isolates from Brazil.^10^ In our analysis, we did not observe any effects of MSL gene deletions on mortality or any other trait (GWAS *P*-values for mortality all MSL genes > 1×10^−2^), consistent with MSL being related to miltefosine resistance, rather than mortality. The lack of association between the MSL locus and mortality is expected for our dataset, as none of the evaluated patients were treated with miltefosine.

We consider it unlikely that the associations between mortality and sex with parasite genotype described here are due to drug resistance. This cohort is not ideal for such as analysis, since patients were treated with a variety of drugs, depending on disease severity (meglumine antimoniate, amphotericin B, liposomal amphotericin B or pentamidine, see **Supplementary Table S3**). Deaths from VL usually happen in the first week of admission. Drug treatments, even when resistance is present, are effective at this point and resistance is noted only weeks after the beginning of the therapy. At present, no clear relationship between treatment failure and susceptibility for these drugs has been demonstrated in Brazil. Preliminary *in vitro* susceptibility analysis with some of the *L. infantum* isolates examined here confirms the lack of association between patient mortality and parasite drug susceptibility (Mayara Lima, unpublished analyses). Host sex is however implicated in VL incidence^38,39^ and increased parasitic load,^40^ but not in host mortality.^4,5^ Therefore, the links of *L. infantum* genome to mortality and sex seems to follow distinct pathogenic pathways.

Our GWAS analysis indicated that a variety of different genetic variants were associated with clinical outcomes (**Table 2**). We discuss the potential implications of each observed SNP/indel and GCNV in **Supplementary Texts B** and **C**, respectively. While we estimate that the parasite genotype has a strong influence on the chance of mortality (*h*^2^ = 0·83 ±0·17, 95% confidence interval 0·66-1·00), and some proportion of this effect appears to be derived from SNPs (**Figure 2**), we did not observe any SNPs that were significantly associated with mortality (**Figure 3**). This is possibly due to mortality being influenced by multiple alleles, each with small effects. While the significance values in our GWAS analysis appear fairly modest compared to GWAS analysis of human genetic variants that affect VL susceptibility,^17^ several considerations need to be taken into account. First, we have a smaller sample size

(*n*=109), so we cannot expect such strongly significant associations from a GWAS. Our results remain significant after permutation, because the *Leishmania* genome is smaller than human/dog genomes, which reduces the burden of multiple test correction. Nevertheless, some of our associations are very strong, particularly when covariate analysis is carried out. The gene LINF_340051500 was supported by a robust *P*-value of 6·4×10^−9^, when the Kala-Cal** was used as a covariate. Similarly, the genes LINF_270006900 and LINF_330042400 were supported when each trait besides Kala-Cal* and ** was simultaneously used as covariate (*P*-values of respectively 5·9×10^−7^ and 1·19×10^−6^, **Supplementary Table S9**), and LINF_270006900 was also supported in all cases when each other trait was individually used as covariates (*P*-values ranging from 8·08×10^−7^ to 1·25×10^−8^; **Supplementary Table S9**).

Many of the variants associated with clinical outcomes defy mechanistic explanations. This is unsurprising given the poor state of basic knowledge about *Leishmania*-host interactions. Intergenic copy variants are particularly challenging to explain mechanistically, as the function(s) of intergenic regions in *Leishmania* genomes are not well understood. However, intergenic regions in the human genome and yeast genomes are frequently associated with phenotypes, consistent with known functional elements within intergenic regions of these species.^34,41^

We found no associations between chromosome somy (CCNVs) and phenotypic traits. This is in accordance with previous work suggesting that karyotypic levels change rapidly between insect and mammalian hosts, and also when parasites are cultured while GCNVs are drivers of long-term adaptation in the field.^31^ As the DNA from the parasites were obtained after only two passages, and significant changes in parasite virulence due to culturing are only observed after 20 to 30 passages^42^ we believe that SNPs and GCNV alterations due to culturing would be minimal.

Furthermore, even if small (SNP/indel) variations are altered within cultured parasites; such variations were previously associated with a reduction in virulence, which is restored with passages in mice.^42,43^ Hence, genetic changes with parasite culture would be expected to *reduce* the strength of our GWAS/heritability associations between parasite genotype and clinical outcomes, rather than create artifactual ones. The strong relationships between parasite genotype and clinical factors observed in the present work indicates that the genome data obtained from cultured parasites relates meaningfully to outcomes.

We envisage several implications of our results. Most importantly, the strong influence of parasite genetic variation on mortality indicates that *L. infantum* isolates vary in their virulence, so the risk of mortality depends at least in part on the isolate that causes the infection. Variation in parasite virulence within Brazilian *L. infantum* isolates may explain the range of clinical VL symptoms, from symptomatic infections,^44^ to rare occurrences of highly-virulent infections that result in high fatality rate for VL in Brazil.^45^ While it may be possible to use parasite genotypes as early predictors of severe infections, this is unlikely to be economically viable, or practically feasible at present, given the complexity of the genotypes associated with mortality and the requirement for parasite culture. It is more practical to use clinical data for risk assessment which are well-powered and feasible in resource-poor settings^5^ (see **Supplementary Figure 14**). However, studies of this type will lead to an enhanced understanding of the molecular mechanisms that lead to severe VL disease. The pathway to this understanding will be a combination of assessing genetic markers, the parasite genes that are implicated, and the host genes enrolled in the immune responses that the different alleles elicit. As many *Leishmania* genes lack detailed gene functional annotations, laboratory analysis will be required to give mechanistic insights.

## Abbreviations

VL: visceral leishmaniasis
GWAS: genome-wide association study
CNV: copy number variant
GCNV: gene copy number variant
IGCNV: intergenic copy number variant
CCNV: chromosome copy number variant
REML: restricted maximum likelihood
LD: linkage disequilibrium
PCA: principal component analysis
RDC: read depth coverage
CV: coefficient of variation
SNP: single nucleotide polymorphism
SNV: small nucleotide variant
HLA: human leukocyte antigen
CL: cutaneous leishmaniasis
SD: standard deviation
ML: maximum likelihood

## Author contributions

CAG - performed genomics and analysis, contributed to writing of the manuscript, and verified the underlying data.

KSSC - performed isolate culture, DNA extraction for 65 *L. infantum* isolates, obtained genomic sequences.

MISL - performed isolate culture and DNA extraction for 14 Brazil samples, phenotypic and genetics analysis, verified the underlying data, and contributed to writing of the manuscript.

VCS - performed isolate culture and DNA extraction for 30 Brazil samples, obtained genomic sequences, and took care of the parasite collection.

JLRC - performed ploidy and CNV analysis, contributed to writing of the manuscript, and verified the underlying data.

MJB - performed ploidy and CNV analysis, contributed to writing of the manuscript.

SJF - contributed to the variant calling pipeline.

CMPSA - performed sample collection, clinical diagnosis, treatment and follow up of patients.

DLC - contributed to collecting and gathering patient clinical data.

JCM - obtained funding, contributed to study design, and contributed to writing of the manuscript.

DS - provided support and advice with heritability and GWAS.

DCJ - obtained funding, developed study design, performed analysis, verified the underlying data and contributed to writing of the manuscript.

CHNC - obtained funding, conceived the study, verified the underlying data, and contributed to study design, collection of patient clinical data and isolates and to writing of the manuscript.

## Full funding details

CAG was supported by MRC Newton as a component of the UK:Brazil Joint Centre Partnership in Leishmaniasis to JCM (MR/S019472/1). JLRC and DCJ are supported by a MRC New Investigator Research Grant to DCJ (MR/T016019/1). SJF was supported by a Wellcome Trust Seed Award in Science to DCJ (208965/Z/17/Z).

MISL and CMPSA are supported by CAPES - Coordination for the Improvement of Higher Education Personnel (Finance Code 001 and PROCAD-Amazônia 001- 21/2018) and FAPEMA - Foundation for Research and Scientific and Technological Development of Maranhão. CHNC was supported by the National Council of Scientific and Technological Development (CNPq). The Brazilian Ministry of Education supported this research.

## Acknowledgements

This project was undertaken on the Viking Cluster, which is a high performance compute facility provided by the University of York. We are grateful for computational support from the University of York High Performance Computing service, Viking and the Research Computing team. We also thank The Genomics & Bioinformatics Laboratory at the University of York for assistance with genome sequencing. We thank Natan Portella Tropical Disease Institute for providing access to the patients and clinical data. We also thank the Brazilian Legislators that provided funds for the Ministry of Health destined to this study.

## Declaration of interests

The authors state that no conflicts of interest exist.

## Data availability

All **Supplementary Materials**, including tables, are available online at the following FigShare address: https://figshare.com/account/home#/projects/126331.

Genome sequencing data produced here are available from NCBI’s sequencing read archive under the BioProject ID PRJNA781413. All other sequencing data used are listed in **Supplementary Table S1**.

## References

1. Ribeiro, C. J. N. et al. Space-time risk cluster of visceral leishmaniasis in Brazilian endemic region with high social vulnerability: An ecological time series study. PLoS Negl. Trop. Dis. 15, e0009006 (2021).

2. Burza, S., Croft, S. L. & Boelaert, M. Leishmaniasis. Lancet 392, 951–970 (2018).

3. WHO. Control of the leishmaniases WHO technical report series 949. Geneva, Switzerland: World Health Organization (2010).

4. Costa, C. H. N. et al. Is severe visceral leishmaniasis a systemic inflammatory response syndrome? A case control study. Rev. Soc. Bras. Med. Trop. 43, 386– 392 (2010).

5. Costa, D. L. et al. Predicting death from kala-azar: construction, development, and validation of a score set and accompanying software. Rev. Soc. Bras. Med. Trop. 49, 728–740 (2016).

6. Costa, D. L. et al. Serum cytokines associated with severity and complications of kala-azar. Pathog. Glob. Health 107, 78–87 (2013).

7. Porcino, G. N. et al. Evaluation of methods for detection of asymptomatic individuals infected with Leishmania infantum in the state of Piauí, Brazil. PLoS Negl. Trop. Dis. 13, e0007493 (2019).

8. Silva de Lima, U. R., Vanolli, L., Moraes, E. C., Ithamar, J. S. & Azevedo, C. de M. P. E. S. de. Visceral leishmaniasis in Northeast Brazil: What is the impact of HIV on this protozoan infection? PLoS One 14, e0225875 (2019).

9. Alvar, J. et al. Leishmaniasis worldwide and global estimates of its incidence. PLoS One 7, e35671 (2012).

10. Carnielli, J. B. T. et al. A Leishmania infantum genetic marker associated with miltefosine treatment failure for visceral leishmaniasis. EBioMedicine 36, 83–91 (2018).

11. Pan American Health Organization. Leishmaniasis. Epidemiological Report of the Americas, March 2019. Leishmaniasis report;7 (2019).

12. Cota, G., Erber, A. C., Schernhammer, E. & Simões, T. C. Inequalities of visceral leishmaniasis case-fatality in Brazil: A multilevel modeling considering space, time, individual and contextual factors. PLoS Negl. Trop. Dis. 15, e0009567 (2021).

13. Lima, I. D. et al. Leishmania infantum chagasi in northeastern Brazil: asymptomatic infection at the urban perimeter. Am. J. Trop. Med. Hyg. 86, 99– 107 (2012).

14. Silva, L. P. et al. Asymptomatic Leishmania infection in blood donors from a major blood bank in Northeastern Brazil: a cross-sectional study. Rev. Inst. Med. Trop. Sao Paulo 62, e92 (2020).

15. Maia-Elkhoury, A. N. S. et al. Premature deaths by visceral leishmaniasis in Brazil investigated through a cohort study: A challenging opportunity? PLoS Negl. Trop. Dis. 13, e0007841 (2019).

16. Mohamed, H. S., Ibrahim, M. E. & Blackwell, J. M. 4. Genetic susceptibility to visceral leishmaniasis. The genetics of African populations in health and disease. Cambridge University Press, Cambridge 71–85 (2019).

17. Blackwell, J. M., Fakiola, M. & Castellucci, L. C. Human genetics of Leishmania infections. Hum. Genet. 139, 813–819 (2020).

18. Ponte-Sucre, A. et al. Drug resistance and treatment failure in leishmaniasis: A 21st century challenge. PLoS Negl. Trop. Dis. 11, e0006052 (2017).

19. Carvalho, K. S. S. et al. Application of next generation sequencing (NGS) for descriptive analysis of 30 genomes of Leishmania infantum isolates in Middle-North Brazil. Sci. Rep. 10, 12321 (2020).

20. Azevedo, T. S. de, Lorenz, C. & Chiaravalloti-Neto, F. Risk mapping of visceral leishmaniasis in Brazil. Rev. Soc. Bras. Med. Trop. 52, e20190240 (2019).

21. Casadevall, A. & Pirofski, L.-A. Virulence factors and their mechanisms of action: the view from a damage-response framework. J. Water Health 7 Suppl 1, S2–S18 (2009).

22. Moxon, R., Reche, P. A. & Rappuoli, R. Editorial: Reverse Vaccinology. Front. Immunol. 10, 2776 (2019).

23. Grace, C. A. et al. Candidates for Balancing Selection in Leishmania donovani Complex Parasites. Genome Biol. Evol. 13, (2021).

24. Li, H. & Durbin, R. Fast and accurate long-read alignment with Burrows-Wheeler transform. Bioinformatics 26, 589–595 (2010).

25. Li, H. et al. The Sequence Alignment/Map format and SAMtools. Bioinformatics 25, 2078–2079 (2009).

26. Depristo, M. A. et al. A framework for variation discovery and genotyping using next-generation DNA sequencing data. Nat. Genet. 43, 491–498 (2011).

27. Speed, D., Holmes, J. & Balding, D. J. Evaluating and improving heritability models using summary statistics. Nat. Genet. 52, 458–462 (2020).

28. Chang, C. C. et al. Second-generation PLINK: rising to the challenge of larger and richer datasets. Gigascience 4, 7 (2015).

29. Franssen, S. U. et al. Global genome diversity of the Leishmania donovani complex. eLife vol. 9 (2020).

30. Rogers, M. B. et al. Chromosome and gene copy number variation allow major structural change between species and strains of Leishmania. Genome Res. 21, 2129–2142 (2011).

31. Bussotti, G. et al. Leishmania Genome Dynamics during Environmental Adaptation Reveal Strain-Specific Differences in Gene Copy Number Variation, Karyotype Instability, and Telomeric Amplification. MBio 9, (2018).

32. Domagalska, M. A. et al. Genomes of Leishmania parasites directly sequenced from patients with visceral leishmaniasis in the Indian subcontinent. PLoS Negl. Trop. Dis. 13, e0007900 (2019).

33. Power, R. A., Parkhill, J. & de Oliveira, T. Microbial genome-wide association studies: lessons from human GWAS. Nat. Rev. Genet. 18, 41–50 (2017).

34. Jeffares, D. C. et al. The genomic and phenotypic diversity of Schizosaccharomyces pombe. Nat. Genet. 47, 235–241 (2015).

35. Alves, F. et al. Recent Development of Visceral Leishmaniasis Treatments: Successes, Pitfalls, and Perspectives. Clin. Microbiol. Rev. 31, (2018).

36. Imamura, H. et al. Evolutionary genomics of epidemic visceral leishmaniasis in the Indian subcontinent. Elife 5, (2016).

37. Downing, T. et al. Whole genome sequencing of multiple Leishmania donovani clinical isolates provides insights into population structure and mechanisms of drug resistance. Genome Res. 21, 2143–2156 (2011).

38. Sharma, M. C., Gupta, A. K., Saran, R. & Sinha, S. P. The effect of age and sex on incidence of kala-azar. J. Commun. Dis. 22, 277–278 (1990).

39. Gouvêa, M. V., Werneck, G. L., Costa, C. H. N. & de Amorim Carvalho, F. A. Factors associated to Montenegro skin test positivity in Teresina, Brazil. Acta Trop. 104, 99–107 (2007).

40. Zacarias, D. A. et al. Causes and consequences of higher Leishmania infantum burden in patients with kala-azar: a study of 625 patients. Trop. Med. Int. Health 22, 679–687 (2017).

41. Giral, H., Landmesser, U. & Kratzer, A. Into the Wild: GWAS Exploration of Non-coding RNAs. Front Cardiovasc Med 5, 181 (2018).

42. Moreira, D. et al. Impact of continuous axenic cultivation in Leishmania infantum virulence. PLoS Negl. Trop. Dis. 6, e1469 (2012).

43. Ali, K. S., Rees, R. C., Terrell-Nield, C. & Ali, S. A. Virulence loss and amastigote transformation failure determine host cell responses to Leishmania mexicana. Parasite Immunol. 35, 441–456 (2013).

44. Costa, C. H. N. et al. Household structure and urban services: neglected targets in the control of visceral leishmaniasis. Ann. Trop. Med. Parasitol. 99, 229–236 (2005).

45. Ruiz-Postigo, J. A. et al. Global leishmaniasis surveillance: 2019-2020, a baseline for the 2030 roadmap/Surveillance mondiale de la leishmaniose: 2019-2020, une periode de reference pour la feuille de route a l’horizon 2030. Weekly Epidemiological Record 96, 401–420 (2021).

